# MFPLI: A Computational Framework for Assessing Biological Authenticity of Protein-Ligand Interactions Using Molecular Fingerprints and Structural Features

**DOI:** 10.1101/2025.06.19.659858

**Authors:** Hao Zhang, Jiqing Zheng, Haoxiang Li, Shizhang Wan, Guiyu Guan, Bin Li, Lei Liu, Wei He

**Affiliations:** School of Pharmaceutical Sciences, Tsinghua University, Beijing, China; New Cornerstone Science Laboratory, Tsinghua-Peking Joint Center for Life Sciences, Ministry of Education Key Laboratory of Bioorganic Phosphorus Chemistry and Chemical Biology, Center for Synthetic and Systems Biology, Department of Chemistry, Tsinghua University, Beijing, China; Foresight Therapeutics (Hefei) Co., Ltd., Hefei, China

**Keywords:** MFPLI, biological authenticity

## Abstract

Traditional computational drug discovery approaches struggle to accurately evaluate the biological authenticity of protein-ligand binding conformations due to inherent limitations in empirical scoring functions and force field approximations. This study proposes MFPLI – a deep learning framework integrating multimodal physicochemical features to systematically assess the biological authenticity alignment between molecular docking poses and true co-crystal structures. By establishing a continuous surface characterization system for protein-ligand interfaces, we concurrently incorporate geometric curvature features (radius, shape index) and chemical interaction fields (electrostatic potential, hydrogen-bond networks, hydrophobicity gradients). A contrastive learning architecture based on Siamese equivariant graph neural networks was developed to enable discriminative analysis between co-crystal conformations and parameter-perturbed pseudo-conformations generated through inverse docking. The five-channel fusion model demonstrates robust performance on the time-split PoseBuster validation set (AUC=0.91), with predicted Euclidean distance deviation (ΔE) effectively distinguishing native co-crystal conformations from aberrant docking poses in 80% of samples. Notably, 71% of ΔE-negative samples concentrate within the [-0.3, 0] interval, reflecting physical consistency between model predictions and conformational transition processes. This framework establishes a novel paradigm for biological authenticity assessment in virtual screening for computer-aided drug discovery through synergistic modeling of surface topology and interaction chemistry.

## INTRODUCTION

Drug discovery remains a complex and protracted endeavor, with traditional approaches demanding exhaustive experimental validation that proves both costly and inefficient.^1^ Transformative innovations in computational drug discovery—particularly the synergistic integration of Computer-Aided Drug Design (CADD) and Artificial Intelligence-Driven Drug Discovery (AIDD)—are currently redefining pharmaceutical research paradigms. Within this evolving technological landscape, classical molecular docking algorithms exhibit significant limitations in quantitative binding free energy prediction due to their reliance on empirical scoring functions.^2^ In contrast, explicit-solvent molecular mechanics conformation sampling methods^3,4^(e.g., Free Energy Perturbation, FEP) and deep generative molecular design frameworks (e.g., diffusion model-driven architectures) have established a multi-scale computational synergy, elevating lead compound discovery to unprecedented efficiencies.^5-7^ Nevertheless, persistent systematic deviations exist between predicted conformations from these methods and experimentally resolved co-crystal structures, manifesting as geometric inconsistencies in ligand topological matching and elevated prediction variance in binding free energy (ΔG) calculations. These discrepancies originate fundamentally from approximation artifacts inherent in classical force field parameterization and inadequate characterization of molecular mechanical constraints by deep learning models. ^8-10^

The core bottleneck resides in the absence of a biologically interpretable evaluation framework,^11^ which critically impedes accurate identification of genuine co-crystal-aligned binding modes within virtual screening-derived conformation ensembles.^12^ While iterative refinement strategies combining molecular docking and FEP are widely adopted for error correction,^13,14^ inherent theoretical constraints prevent these algorithms from resolving the biological authenticity assessment impasse. Notably, deep learning—and specifically the nonlinear representational capacity of geometric deep learning—presents an important breakthrough pathway.^15-17^ Constructing physically constrained, interpretable neural architectures enables extraction of physiologically relevant conformational features from high-dimensional virtual data, thereby facilitating discrimination of biologically plausible conformations.

Building upon this conceptual foundation, we introduce a Siamese contrastive learning architecture employing equivariant graph neural networks.^18^ This framework advances the development of MFPLI (Molecular Fingerprint based Protein Ligand Interaction)—a deep generative model with enhanced conformational discriminative capability. From a molecular representation standpoint, molecular surface characterization serves as a topological descriptor of protein tertiary structure. By modeling within continuous function space, it simultaneously incorporates geometric curvature features (e.g., radius, shape index) and chemical interaction fields (encompassing electrostatic potentials, hydrophobicity gradients, and hydrogen-bond donor/acceptor distributions). This multimodal feature fusion strategy directly reflects the physicochemical essence of ligand-receptor binding interfaces. Implementing MFPLI for virtual screening assessment preserves the energy-driven paradigm of molecular docking while transcending traditional force field parameterization bottlenecks through machine learning, thereby establishing a novel framework for high-fidelity protein-ligand interaction prediction.

## RESULTS

### Deep Learning-Driven Computational Framework for Topological Feature Fusion at Protein-Ligand Interfaces

Protein co-crystal complexes comprise biomacromolecules bound to small-molecule ligands. Initial three-dimensional surface reconstruction of crystalline complexes employs a van der Waals-based solvent-accessible surface (SAS) representation system (Fig. 1A). Subsequent decomposition into individual components enables separate topological characterization: ligand molecules undergo Delaunay triangulation to generate meshed surface models (Fig. 1B). Each vertex in these models undergoes multi-dimensional feature encoding, integrating two geometric parameters (curvature radius, shape index) and three chemical descriptors (electrostatic field strength, hydrogen-bond donor/acceptor loci, and hydrophobic potential distribution), with detailed parameterization methods provided in the Methodology section.

**Fig. 1.**
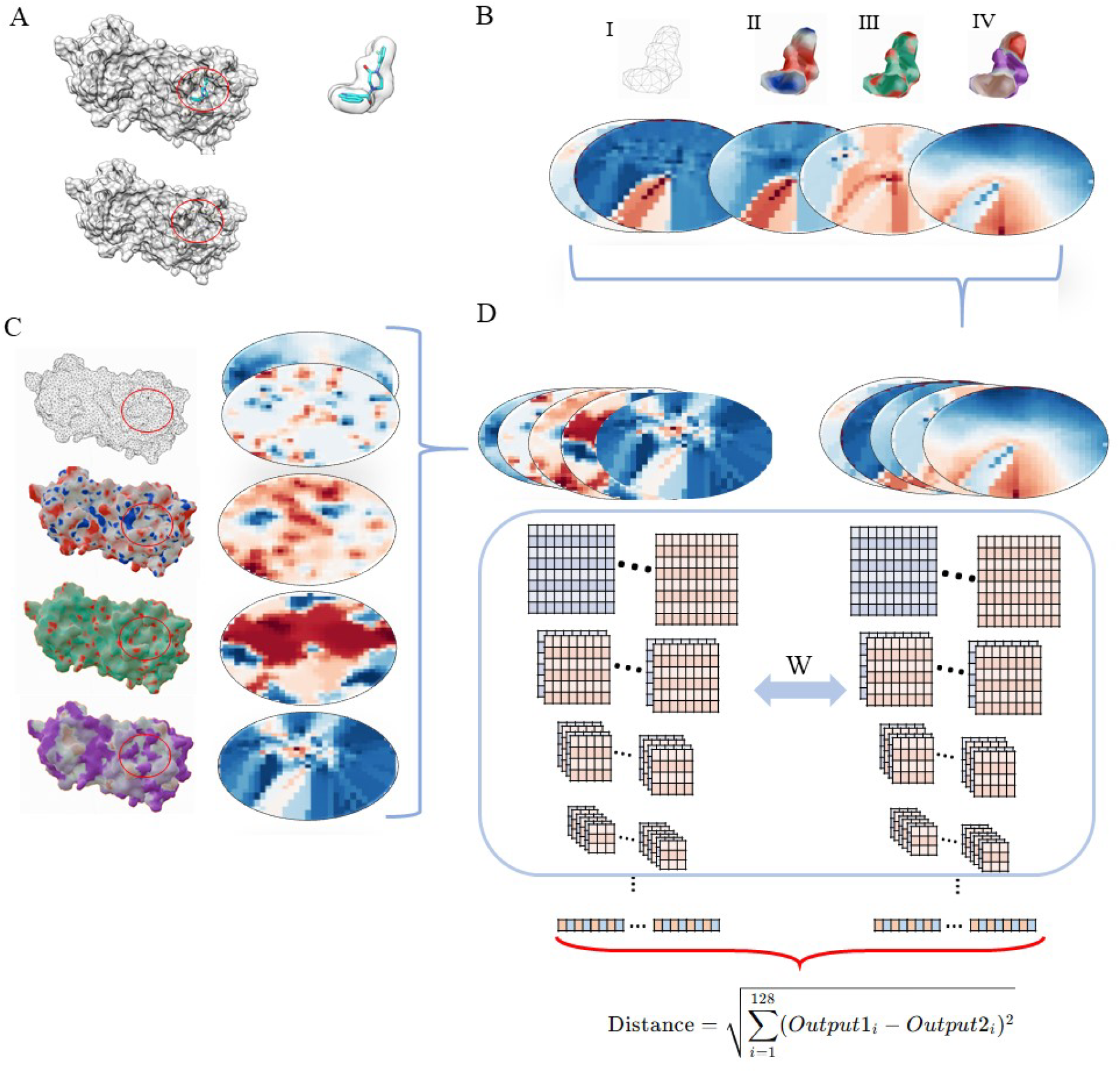
Molecular surface characterization and network architecture. (A) Solvent-accessible surface (SAS) topologies for the receptor (gray) and ligand (cyan) derived from PDB:7GHS cocrystal structure, with spatial dissociation establishing baselines for multimodal feature encoding. (B) Ligand surface processing: (I) Delaunay triangulation; (II) Electrostatic potential mapping; (III) Hydrogen-bond density annotation; (IV) cLogP-based hydrophobic gradient projection. Resulting feature matrices generated via 2D grid projection. (C) Localized receptor surface processing (16Å radius sphere centered at ligand centroid) with identical electrostatic/hydrophobic/H-bond feature encoding. (D) Dual-channel siamese network (blue outline) processing 5-channel inputs: two geometric (curvature, shape index) and three chemical (electrostatics, H-bond, hydrophobicity). Shared-weight ConvNeXt backbone enables scale-invariant feature extraction, with contrastive loss-driven projection to 128D normalized embedding space.

Complementarily, protein interface processing employs a ligand-proximal centroid to define a 12Å-radius spherical region for feature extraction (Fig. 1C), capturing identical quintuple parameter categories (two geometric + three chemical). During feature fusion, ligand and protein descriptors are concatenated as five-channel input tensors (Fig. 1D). A Siamese neural network architecture computes feature-space Euclidean distances to evaluate topological congruence between co-crystal conformations. Crucially, experimentally resolved co-crystal structures exhibit high feature-space similarity, while conformationally aberrant poses demonstrate pronounced deviation.

Model training utilizes ConvNeXt deep residual networks as dual-branch encoders under parameter sharing constraints. Fully connected decoder layers with nonlinear activations reduce dimensionality, ultimately outputting scalar binding compatibility metrics. Optimization employs contrastive loss with dynamic margin adjustment, enhancing intra-class compactness and inter-class separation within the learned latent space.

### Multidimensional Chemical Feature Validation and Conformational Perturbation Strategy

Prior to machine learning implementation, we rigorously validated the theoretical underpinnings of our feature selection strategy through chemical pattern recognition analysis. This a priori assessment determined whether the chosen physicochemical descriptors could capture fundamental co-crystallization principles. Using SARS-CoV-2 main protease complexes as a model system, multidimensional feature visualization demonstrated distinct patterns of molecular recognition: In the electrostatic complementarity dimension (Fig. 2A□), ligand-receptor surface potentials exhibited topological complementarity where ligand-positive regions (blue) spatially coupled with protein-negative zones (red), consistent with electrostatic attraction theory as indicated by yellow and red vector arrows. The hydrogen-bond network analysis (Fig. 2A□) revealed geometrically proximate distributions between donor and acceptor sites, while hydrophobic matching (Fig. 2A□) showed continuous polarity gradients formed between nonpolar ligand surfaces and receptor hydrophobic pockets. These results confirm that the three chemical descriptors follow interpretable physicochemical principles, and their nonlinear combinations contain higher-order correlative features extractable by deep convolutional networks, thereby validating our discriminative feature set.

**Fig. 2.**
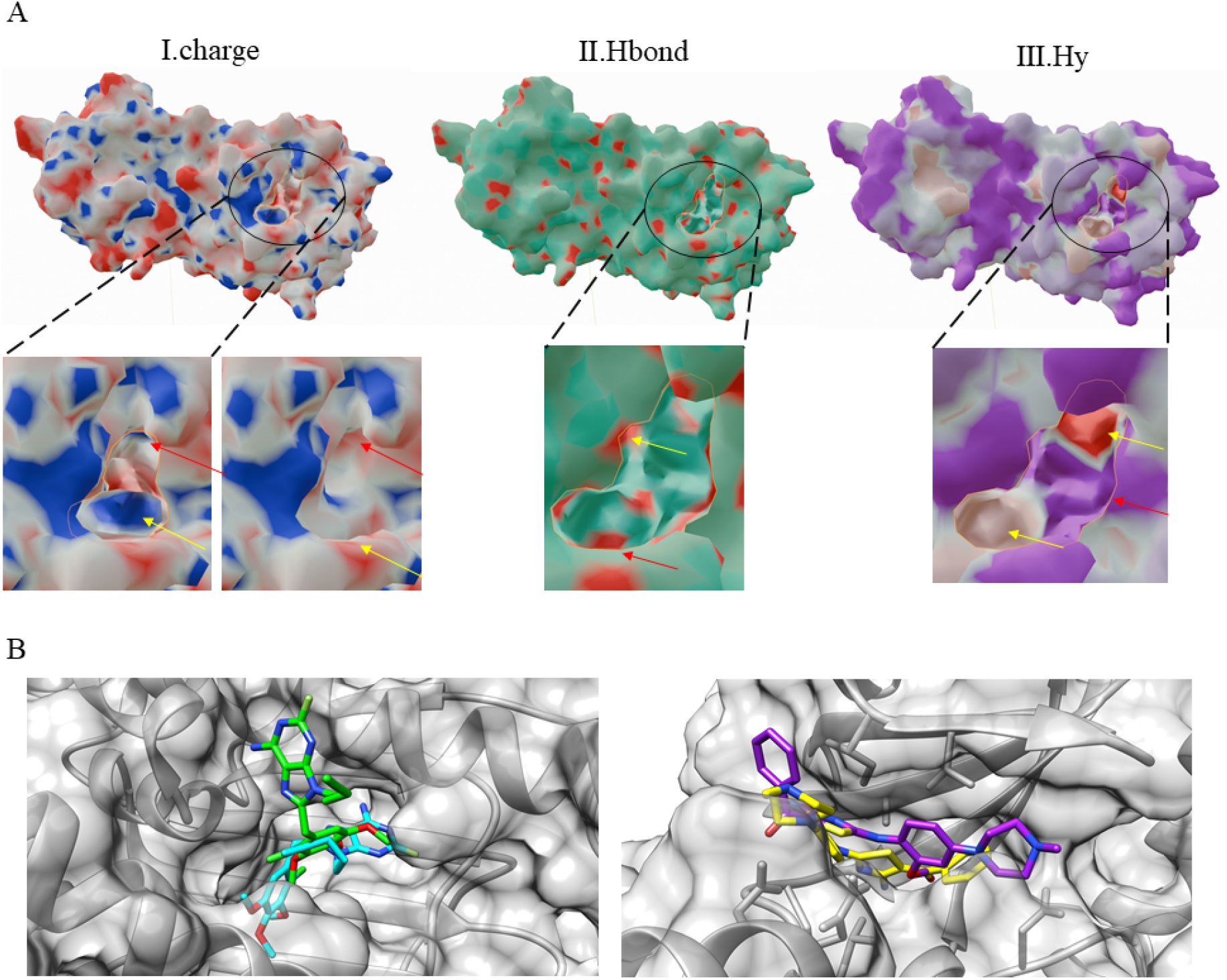
Interface characterization and discriminative performance. (A) PDB:7GHS interface analysis: (I) Spatial complementarity between ligand low-charge (red) and receptor high-charge zones (red arrows); (II) Electrostatic-H-bond synergy in binding regions; (III) Hydrophobic-polar complementarity following “like-dissolves-like” principle. (B) Conformational discrimination: (I) Distinct surface feature mismatch between native ligand (green) and high-RMSD pose (RMSD=7.97Å, cyan) in PDB:6EL5; (II) Pronounced topological deterioration in PDB:6EIQ (native: purple, decoy RMSD=8.92Å: yellow).

Building upon this verified chemical complementarity framework, we implemented an inverse docking strategy to generate conformationally diverse negative samples. Through systematic perturbation of Autodock Vina parameters—reducing weight_gauss1 to 0.015 while inverting weight hydrophobic and weight hydrogen to 0.2 and 0.5 respectively—we performed comprehensive conformational space sampling of co-crystal complexes. As evidenced in Fig. 2B, this parametrically modified docking protocol consistently generated two classes of non-native binding modes: spatially displaced poses with ligand binding site offsets exceeding 5Å, and rotationally inverted poses exhibiting 180° ligand orientation reversals. This controlled perturbation strategy effectively expanded negative sample diversity, enabling subsequent deep learning models to discern co-crystal-specific interaction fingerprints with enhanced discrimination capability.

### Deciphering Synergistic Interplay of Chemical Determinants and Physical Topological Constraints

To elucidate the cooperative mechanism underlying fused feature modeling, we implemented feature decomposition analysis evaluating the relative contributions of chemical and physical descriptors to co-crystal discrimination. Our multidimensional architecture integrates three chemical interaction channels (electrostatic potential, hydrogen-bond density, hydrophobic gradients) with two physical topological descriptors (surface curvature radius, shape index). As quantified in Fig. 3B, the five-channel fused model demonstrated robust discrimination performance (AUC=0.91), with Euclidean distance distributions revealing significant separability between positive and negative samples. Crucially, exclusive use of dual-channel physical topology descriptors (Fig. 3C) achieved marginally higher AUC (0.95), yet exhibited constrained feature space dispersion—indicating overreliance on local topological similarity at the expense of expressive capability. Conversely, the triple-channel chemical interaction model (Fig. 3D) maintained high AUC (0.92) while delivering superior feature separability. This alignment with the chemically dominant recognition patterns established in Fig. 2A demonstrates that chemical descriptors achieve an optimal balance between mechanistic interpretability and predictive efficacy through multidimensional interaction integration.

**Fig. 3.**
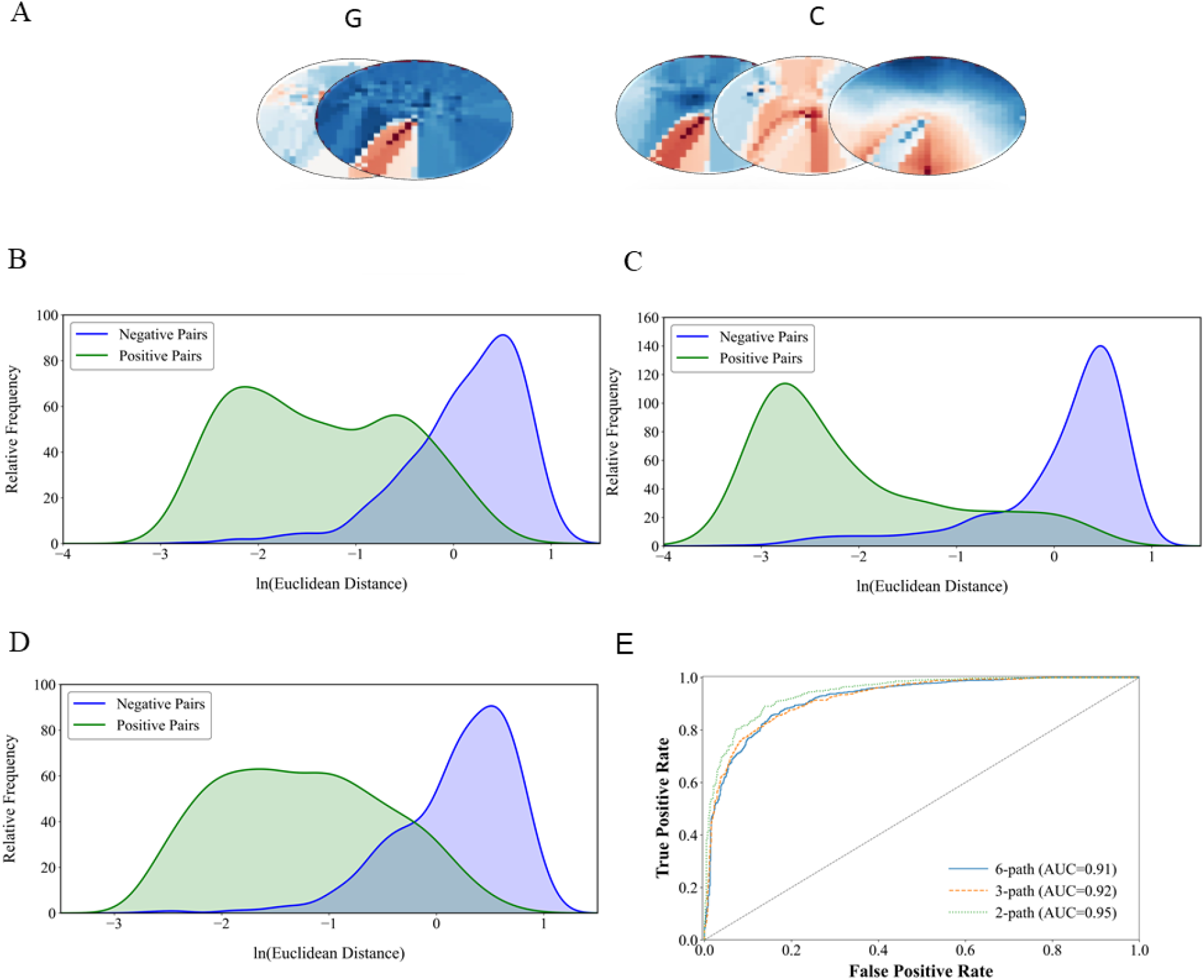
Ablation study of feature channel contributions. (A) Input channel schema: geometric (G: 2 channels) vs. chemical (C: 3 channels). (B-D) Euclidean distance distributions for: (B) Full G+C model, (C) G-only, (D) C-only architectures. (E) Convergence dynamics: G+C (AUC=0.91 at 80 epochs), G-only (AUC=0.95 at 50 epochs), C-only (AUC=0.92 at 70 epochs), revealing accelerated geometric feature optimization.

Convergence analysis (Fig. 4A) revealed fundamental operational differences: the physical topology model reached peak performance (AUC=0.95) at 50 training epochs, while the chemical feature model required 70 epochs for optimal feature space refinement. The fused architecture demonstrated the longest convergence trajectory (80 epochs). This phenomenon highlights two key mechanisms: Physical descriptors efficiently capture geometric constraints despite limited information dimensionality, whereas chemical features necessitate extended training to resolve high-order nonlinear relationships governing molecular recognition. Notably, the fused model’s ultimate discrimination performance did not proportionally increase with feature dimensionality due to gradient conflict arising from nonlinear cross-modal coupling. This multimodal analysis substantiates our selection of the five-channel architecture—chemical descriptors provide dominant recognition signals while physical constraints deliver complementary spatial regularization, thus critically optimizing conformational sampling efficiency in virtual screening applications.

**Fig. 4.**
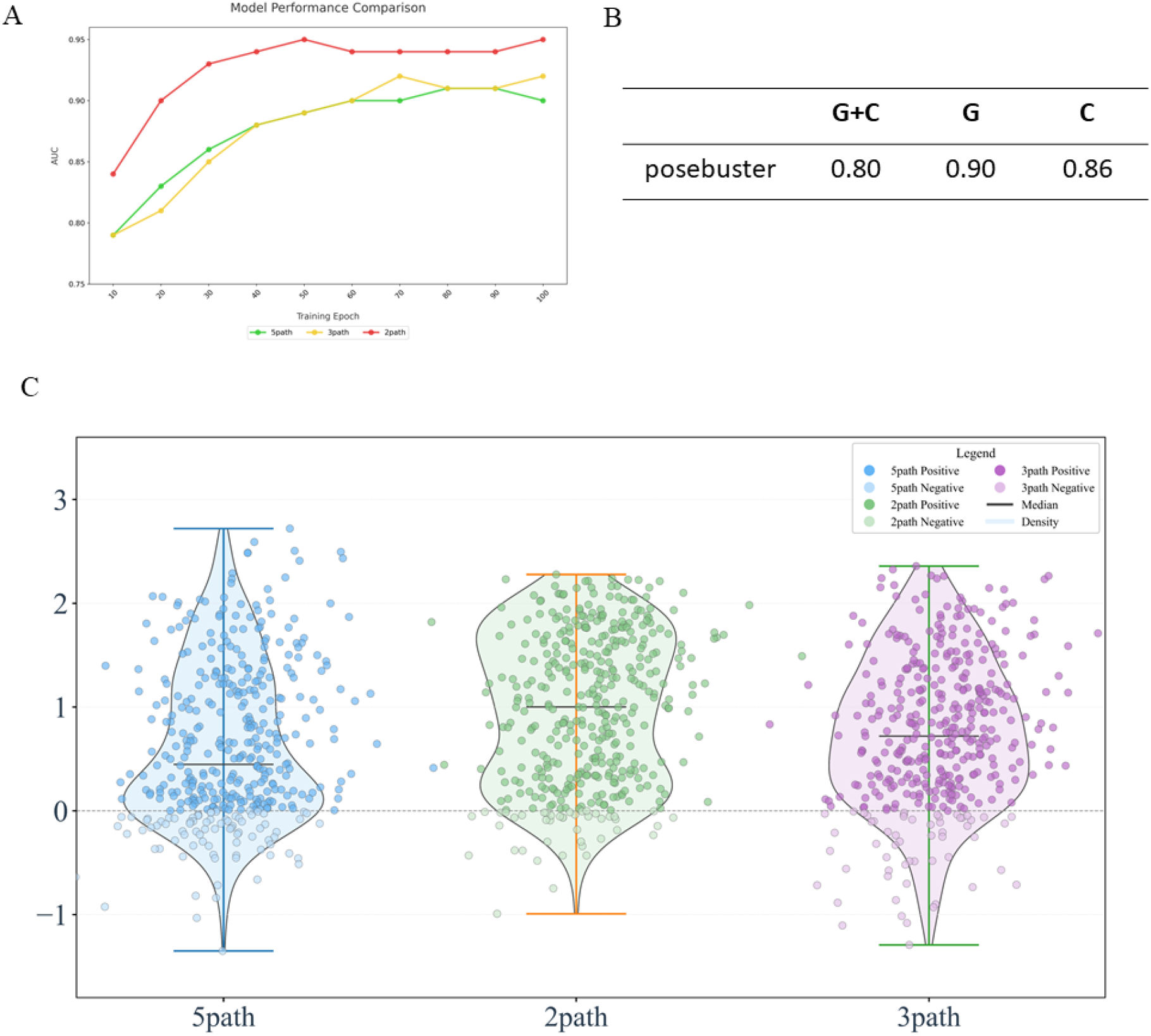
Discriminative performance evaluation of multi-channel architectures. (A) Five-channel fusion (G+C) achieved peak AUC (0.91) at 80 epochs, while dual-channel physical (G) and triple-channel chemical (C) models converced faster at 50 (AUC=0.95) and 70 epochs (AUC=0.92), respectively. (B) On PoseBuster, G+C showed balanced accuracy (80%). The G model yielded higher accuracy (90%) but anomalous error distribution, whereas C declined to 86%. (C) ΔE distributions show: G+C: 80% true positives (ΔE>0; dark) with moderate dispersion (broad density contour) G: 90% ΔE>0 with compact unimodal distribution (high peak) C: 86% ΔE>0 exhibiting elongated tails and higher dispersion

### Discriminative Efficacy and Generalization Superiority of Multi-Feature Fusion Models

To systematically evaluate model generalization in unexplored conformational space, we implemented a time-split validation strategy using the PoseBuster dataset (non-redundant crystal structures released post-2020) as an external benchmark, ensuring strict temporal independence from the PDBbind V2020 training set. Comparative analysis of multimodal architectures (five-channel physicochemical fusion, dual-channel physical topology, triple-channel chemical features) revealed fundamental principles governing co-crystal discrimination robustness. Quantitative analysis of output distributions (Fig. 4C) demonstrated that Euclidean distance deviations (ΔE = D_negative_−D_positive_) for the five-channel model exhibited significantly positive skewness within the PoseBuster test set. Critically, 80% of samples displayed ΔE > 0, confirming the model’s efficacy in distinguishing native co-crystal conformations from aberrant docking poses through superior prediction of ligand-protein spatial arrangements. This discriminative sensitivity originates from the multimodal physicochemical representation, achieving 80% high-confidence accuracy in conformation classification.

Notably, negative-deviation samples (ΔE < 0) demonstrated biologically meaningful distribution patterns: 71% concentrated within the [-0.3, 0] interval. This continuous deviation spectrum correlates strongly with physical principles governing gradual conformational transitions, confirming the model’s prediction errors remain bounded within physically plausible limits without systemic drift. The dual-channel physical topology model achieved higher nominal accuracy (90%, Fig. 4B), with misclassified poses universally exhibiting ΔE approximations approaching zero—demonstrating exceptional robustness in shape-based discrimination. Conversely, the chemical complementarity model showed reduced accuracy (86%, Fig. 4B), where negative ΔE samples displayed divergent distribution patterns indicative of feature representation blind spots in complex chemical environments.

Empirical evidence establishes the five-channel fusion model’s unique advantages in informational completeness: While maintaining 80% high discriminative accuracy, it demonstrates substantially more stable ΔE distribution characteristics than unimodal frameworks. Particularly significant is its optimized balance between positive deviation predominance and controlled dispersion of negative deviations. Although the physical model excels in singular accuracy metrics, multidimensional evaluation encompassing robustness, interpretability, and domain adaptability confirms that fused-feature architectures provide superior solutions for discriminating complex pharmaceutical co-crystal systems.

## CONCLUSIONS

This study establishes a deep learning framework that integrates chemical complementarity and physical topological constraints at protein-ligand interfaces to decipher the biological authenticity of native co-crystal conformations. Chemical features—including electrostatic potentials, hydrogen-bond networks, and hydrophobicity gradients—directly encode experimentally validated interaction principles, capturing the essential recognition signatures of native binding modes. Conversely, physical topological parameters (curvature radii and shape indices) provide spatially constrained representations of surface geometry to guide conformational discrimination. The five-channel fusion architecture demonstrated synergistic power through nonlinear integration of physicochemical determinants, where the predicted Euclidean distance deviation metric distinguished native from non-native conformations in 80% of PoseBuster test cases. The ΔE distribution profile confirms strict adherence to physicochemical principles governing co-crystal complexes, with the parametric perturbation strategy applied during inverse docking further enhancing recognition of biology-anchored interaction fingerprints. This methodology thus pioneers a biologically authentic conformation assessment paradigm for virtual screening in pharmaceutical discovery.

## METHODS

### Implementation of Molecular Surface Characterization

Small Molecule Surface Parameterization: A multiscale integrative framework was developed to quantify ligand surface properties, combining molecular topology analysis, quantum chemical computations, and continuum solvation models. Protonated conformations were initially parsed from PDB files using the RDKit cheminformatics platform (MolFromPDBFile function). To overcome limitations in conventional surface characterization, a fragment-based Crippen approximation was employed, whereby atomic-level hydrophobic contribution tensors were resolved via the GetAtomContribs function, thereby enhancing hydrophobic potential accuracy For charge distribution modeling, partial atomic charges were computed using OpenBabel 3.0 with the MMFF94s forcefield (--partialcharge mmff94 flag), incorporating optimized Born-Oppenheimer approximation criteria. Missing charge data were interpolated to grid vertices via KDTree nearest-neighbor searches using atomic hydrophobicity values.

Precise electrostatic potential field generation leveraged a multiscale continuum coupling algorithm. Molecular topologies were reconstructed in Biopython, followed by generation of MSMS-compatible files using customized atomic radii parameters. During surface tessellation, optimized probe radii and grid density parameters concurrently ensured geometric fidelity of the solvent-accessible surface while maintaining computational tractability. APBS solver integration with adaptive mesh refinement techniques substantially improved numerical stability in solving the nonlinear Poisson-Boltzmann equation through theoretical optimization of dielectric constants and innovative spatial discretization. This hierarchical architecture advances theoretical frameworks for molecular surface characterization and provides novel methodological support for predicting intermolecular interactions in drug design.

Protein Surface Parameterization: A multi-tool computational framework (primarily utilizing MaSIF algorithms for feature extraction) unified structure parsing, electrostatic calculations, and physicochemical property mapping. PDB topologies were processed using Biopython’s PDBParser, with protonation states optimized via pdb2pqr (--ff=parse forcefield) for automated polar hydrogen addition. Hydrogen-bonding potentials were calculated using geometric constraint models.

Hydrophobicity was directly mapped using Kyte-Doolittle residue scales. Amino acid types were resolved via vertex identifiers, enabling rapid residue-to-scale value lookup. This approach obviates fragmented atomic contribution interpolation, enabling efficient residue-level property transfer.

Electrostatic potentials were computed using an APBS workflow: input files generated via pdb2pqr were processed by APBS to solve the Poisson-Boltzmann equation, with results interpolated onto surface vertices using multivalue.

Dynamic atom typing was implemented for hydrogen-bond prediction: His ND1/NE2 protonation states determined acceptor eligibility (detecting absent HD1/HE2 atoms), while backbone hydrogen bonds were excluded via NeighborSearch spatial queries. Charge interpolation employed a KDTree fourth-nearest-neighbor weighted scheme, assigning values inversely proportional to squared distances to preserve charge conservation during surface reconstruction.

This integrated framework—combining pdb2pqr-APBS-multivalue continuum modeling with geometrically constrained hydrogen-bond potentials and direct residue hydrophobicity mapping—delivers multidimensional surface characterization data (electrostatics, hydrogen-bonding, hydrophobicity) for robust molecular interaction prediction.

### Model Architecture and Training Configuration

The model employs a siamese network architecture built upon a modified ConvNeXt backbone, leveraging weight-sharing mechanisms to achieve efficient biomolecular similarity metrics. This framework utilizes dual parallel processing pathways, where feature extraction layers inherit from ImageNet-pre-trained ConvNeXt models (supporting tiny/small/base/large parameter scales). Parameter sharing ensures feature space consistency across input pairs. Notably, the classification head of the original model was replaced, retaining only its hierarchical convolutional feature extractor. To accommodate multimodal biomolecular input, the first convolutional layer was extended from 3 to 6 channels, formalized as:

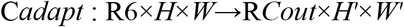

Feature transformation proceeds through three sequential stages: hierarchical feature extraction via ConvNeXt blocks, dimensionality reduction to 1×1×768 embeddings using adaptive average pooling, and projection into 128-dimensional normalized space through the operation F*proj* =σ(W2⋅δ(W1⋅F*pool*+b1)+b2). where σ denotes the Sigmoid activation, δ represents the ReLU nonlinearity, and W1,2, b1,2 correspond to fully-connected layer parameters.

Input data processing involved standardized restructuring of 11-channel .npy files into two distinct six-channel tensors: protein features *X*_protein_∈R6×*H*×*W* (Channels 1-5 + Channel 11) and ligand features *X*_ligand_∈R6×*H*×*W* (Channels 5-11). Spatial standardization to 224×224 resolution was achieved via bicubic interpolation, followed by channel-wise normalization

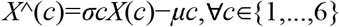

with preset parameters μ*c*=0.5 and σ*c*=0.5.

During inference, the siamese network generates normalized 128-dimensional feature vectors (*fi*,*fj* ) for input pairs (*Xi*,*Xj*), with pairwise distance quantified using the *L*2-norm metric *d*(*fi*,*fj*)=∥*fi*−*fj*∥ 2. This distance measure couples with a dynamic-margin contrastive loss to drive discriminative feature learning through backpropagation. For computational efficiency, mixed-precision training (FP16) was deployed alongside gradient clipping (γ=5.0) to ensure stability.

The framework was implemented in PyTorch with key hyperparameters configured as follows: Adam optimizer (learning rate=10^−6^, weight decay=0.001), mixed-precision computation (FP16), gradient clipping threshold (max|**g**|_2_=5.0), and fixed random seed (torch.manual_seed(42)) with version-controlled dependencies for full reproducibility. All experiments utilized NVIDIA RTX 5090D accelerators.

### Software Implementation

Protein structure parsing and processing employed Biopython’s PDB module.^19^ Ligand format conversion and charge calculations utilized Open Babel,^20^ while molecular surface mesh generation leveraged PyMesh. Molecular surface feature extraction and interaction fingerprint computation integrated the MaSIF framework.^21^ Spatial neighbor searches implemented scikit-learn’s KDTree, with molecular surface topology networks processed using NetworkX. Three-dimensional visualizations were generated with Plotly.

Molecular docking simulations were performed with AutoDock Vina,^22^ and ligand chemical feature analysis utilized RDKit. Numerical computations relied on NumPy and SciPy,^23^ with protein secondary structure calculations implemented via DSSP.^24^ Decoy conformation generation employed Vina.

### Data Sources

Protein three-dimensional structures were retrieved in the standardized Protein Data Bank (PDB) format exclusively from the official RCSB PDB platform (https://www.rcsb.org).^25^ The experimental dataset was constructed using protein-ligand complexes curated from the PDBbind database (v2020 release),^26^ which provides rigorously validated biomolecular benchmarks through dual-criterion selection: binding affinity measurements and crystallographic quality assessment.

Negative conformations for molecular docking were generated by defining a cubic sampling space of 25×25×25 Å^3^ centered on each cocrystalized ligand. Docking parameter optimization included adjusted scoring function weights: weight_hydrogen = 0.5, weight_hydrophobic = 0.2, weight_gauss1 = 0.015.

## Code Availability

All computational workflows were implemented in Python, with neural network architectures developed using PyTorch. The full codebase supporting this study, including experimental reproduction scripts, will be made available on GitHub upon publication.^27,28^

## STATEMENTS OF ETHICAL APPROVAL

Not applicable.

## DECLARATION OF INTEREST STATEMENT

The authors declare no competing financial interest.

## ACKNOWLEDGMENTS

The authors acknowledge financial support from the Beijing Natural Science Foundation (Grant No. Z230020) and the National Natural Science Foundation of China (Grant No. T2488301). We thank Profs. Jun Xu (Sun Yat-sen University) and Boxue Tian (Tsinghua University) for helpful discussion.

## Notes

### Competing Interest Statement

The authors have declared no competing interest.

